# Collagen fibre organisation revealed with computational scattered light imaging and polarimetric second harmonic generation microscopy

**DOI:** 10.1101/2025.07.04.663123

**Authors:** Loes Ettema, Viktoras Mažeika, Mehdi Alizadeh, Hamed Abbasi, Virginijus Barzda, Miriam Menzel

## Abstract

Collagen forms dense fibre networks in the human body, with the organisation directly influencing tissue mechanics and function in health and disease. A good understanding of this relation requires proper imaging techniques for visualising the dense collagen network. Previously, computational scattered light imaging was employed as a fast and easy-to-implement technique to retrieve the orientation of multi-directional collagen fibres, but results were not yet validated quantitatively. In this study, we validate the in-plane orientations of collagen fibres determined with computational scattered light imaging by performing comparative measurements with polarimetric second harmonic generation microscopy on rat tendon and bone sections. For rat tendon, sections with and without hematoxylin-and-eosin staining, folded tendon layers, and obliquely cut sections were investigated. Similar fibre orientations were obtained with both techniques in both tissues, with the highest degree of similarity found for in-plane, unidirectional fibres in the tendon sections. The techniques were able to retrieve the orientations of multi-directional crossing collagen fibres in folded rat tendon layers, and results were found to be unaffected by staining. While polarimetric second harmonic generation microscopy provides high resolution and ultrastructural information on collagen, computational scattered light imaging provides large field of view measurements with micrometre resolution.

## Introduction

Numerous types of tissues consist of collagen, with collagen type I being the most prevalent protein in the human body^1–4^. Collagen type I is a fibrillar protein and has a triple helical structure^2–6^. These helices assemble into fibrils, eventually forming fibres with diameters ranging from 10 to 50 μm ^7^. Within tissues, collagen fibres fulfil an important role in providing the desired mechanical properties, such as strength, flexibility, and toughness^1–6^. These properties are heavily dependent on the hierarchical structure of collagen within the tissue^1,4,8–11^. In skin, for example, collagen fibres form interwoven dense networks that provide both flexibility and mechanical strength^4,10,12^. In tendon, on the other hand, the fibres are oriented along the tendon axis, which facilitates the absorption of energy and conduction of tensile forces^1,4,5^.

If the organisation of the fibres does not support the required properties of the tissue, malfunctioning of the tissue may occur. This is, for example, seen in genetic diseases that impair the organisation of hierarchical structures^1,3,4^, after skin burn, in which heat initiates a process that causes collagen molecules to denature^13,14^, and after myocardial infarction, a condition that damages the collagen network^15^. Changes in the organisation and orientation of collagen fibres have also been identified in cancer studies. In breast and gastric cancer, it was shown that oriented collagen fibres in the extracellular matrix (ECM) direct tumour cell intravasation^16^. During metastasis, collagen fibres are reorganised, which changes their alignment from parallel to perpendicular with respect to the tumour boundaries and opens up passages for the migration of tumour cells into surrounding tissue^17–19^. This change in fibre organisation could potentially serve as a new and promising prognostic risk factor for regional or distant metastasis^20–22^.

Given the direct relation between collagen network structure and tissue functioning, both in the unaffected and the diseased state, visualising the intricate fibre network with high resolution is of importance to obtain a better understanding of this relation. Preferably, the tissue should be investigated in its native form, omitting labour-intensive and time-consuming sample preparation procedures, including tissue staining. One label-free technique that has extensively been used in the field of collagen research is second harmonic generation (SHG) microscopy. Due to its high second-order nonlinear optical susceptibility, collagen induces a strong SHG signal and is therefore suitable for being studied with SHG microscopy. The SHG process is highly sensitive to both the polarisation state of the fundamental light and the sample structure in the focal volume, and polarimetric SHG (pSHG) therefore provides additional ultrastructural and orientation information^15,23^, enabling a more detailed characterisation of collagen fibres. In cancer research, the above-mentioned change to the ECM has also been observed with both SHG and pSHG microscopy^22,24–27^. However, although pSHG is a well-established technique in the field, it suffers from a relatively long acquisition time when ultrastructural information is deduced (images need to be acquired for different polarisation states) and a small field of view (FoV) at submicron resolution^10^.

Computational scattered light imaging (ComSLI) could be an alternative, fast, and easy-to-implement technique for visualising collagen fibre networks in biological samples with micrometre resolution and over large fields of view. Requiring only a light emitting diode (LED) light source and camera, it exploits the scattering of visible light by directed structures in order to retrieve their orientations^28,29^. Previous studies on brain tissue slices have shown the utility of ComSLI for retrieving multi-directional nerve fibre pathways with micrometre resolution, also in regions with highly interwoven and crossing fibres. The determined fibre orientations were validated by performing simulation studies and comparing the measurement results with other techniques, such as diffusion magnetic resonance imaging (dMRI), polarised light imaging (PLI), and small-angle X-ray scattering (SAXS)^28–31^. Recently, ComSLI has also been applied to non-brain tissue and differently prepared samples, revealing orientations of collagen, elastin, and muscle fibres^32^. This suggests that ComSLI can be used for applications outside neuroscience and become a useful tool for studying collagen networks and collagen-related diseases. However, while previous results on brain tissues have been validated in multiple ways, a comprehensive validation on non-brain tissues is still missing.

Here, we perform for the first time a comprehensive comparison between collagen networks reconstructed by ComSLI and pSHG in order to validate the utility of ComSLI for retrieving collagen fibre orientations. We focus on the principles behind both techniques for retrieving in-plane orientations of both unidirectional fibres and multi-directional crossing fibres, the utility of ComSLI, and the challenges that still lie ahead. To this end, the same regions of interest were measured with both techniques in rat tail tendon and rat bone sections; both tissues in which collagen type I is the most dominant protein^33,34^. Processing of the data yielded the collagen fibre orientations, which were subsequently compared in a quantitative, pixel-wise manner. We show that both techniques retrieve similar orientations for unidirectional fibre bundles as well as for crossing fibre bundles, and that orientation results are mostly independent of H&E tissue staining and out-of-plane fibre tilt angles. While pSHG is the current state-of-the-art technique in collagen research and provides determination of high-resolution ultrastructural parameters including fibre orientations and molecular susceptibility tensor component ratios^35,36^, ComSLI enables retrieval of fibre orientations with large fields of view applicable for whole-slide imaging, and has been shown to reliably retrieve orientations of multiple crossing fibres.

## Results

For all samples, measurements were first performed with ComSLI (2.87 μm pixel size, 3.91-4.38 μm lateral resolution), followed by pSHG measurements (2.37 μm pixel size, 4.6 μm lateral resolution) of multiple regions of interest. Orientations were calculated for both techniques independently, after which the upscaled ComSLI data were registered onto the pSHG reference data and the orientations were compared (see Methods for more details). First, we investigated the differences between both techniques in the way they retrieve unidirectional as well as multi-directional crossing fibre orientations, using a sample with unstained rat tail tendon sections. Subsequently, we compared their performances on retrieving unidirectional collagen fibres in hematoxylin and eosin (H&E) stained and eosin-stained tendon samples to study the influence of the sample preparation process on the computed fibre orientations. Afterwards, we analysed samples with out-of-plane tilted fibres using rat tail tendon sections that were obliquely cut with respect to the tendon axis. Finally, we studied the performance of both techniques on a rat bone section, which has a more complex composition of collagen fibres than tendon.

### Extracting in-plane collagen fibre orientations from an unstained rat tendon sample

When illuminating an oriented in-plane structure under normal incidence, scattered light will be projected mainly in the direction perpendicular to the orientation of the structure due to diffraction (Fig. 1a). In ComSLI, it is this principle that is being used, however, measurements are performed in the reversed way: instead of illuminating the sample under normal incidence, light is emitted from multiple azimuth and polar angles using an LED display. The transmitted scattered light is subsequently captured for multiple LEDs on the display, i.e., for multiple illumination angles (Fig. 1b). This results in an image series (Fig. 1c), in which for each pixel within the field of view a scattering pattern can be created by analysing the pixel intensity along the image series. Figure 1d shows the average scattering map of the region indicated by the orange rectangle in Fig. 1c and the scattering patterns for two pixels: one containing a unidirectional bundle of collagen fibres, and one originating from a location where the tendon is folded, therefore resulting in several fibres crossing each other. In these patterns, lines of high scattering intensity indicate the orientation of the fibre bundle, which is, as explained above, perpendicular to this line (see dashed coloured lines). In order to reduce measurement time, measurements can be performed using only one polar angle and a smaller number of azimuth angles, i.e., instead of acquiring the entire scattering pattern, data is obtained along a ring of the pattern (red dotted circles in Fig. 1d) in 15° steps. Plotting the measured intensity against the azimuth angle results in a discretised line profile, in which the mid-position of the peak pairs can be used for determining the fibre orientations (Fig. 1e). Knowing the fibre orientations, each pixel can be assigned an orientation-specific colour according to a colour wheel (Fig. 1f, top right), forming a so-called fibre orientation map (FOM; Fig. 1f, left). In the FOM, each pixel is divided into 2×2 subpixels. If one pixel contains multi-directional crossing fibres, multiple colours will indicate the corresponding fibre orientations (two colours for two different fibre orientations, and three colours plus a black subpixel for three different fibre orientations, see Fig. 1f, bottom right).

**Fig. 1.**
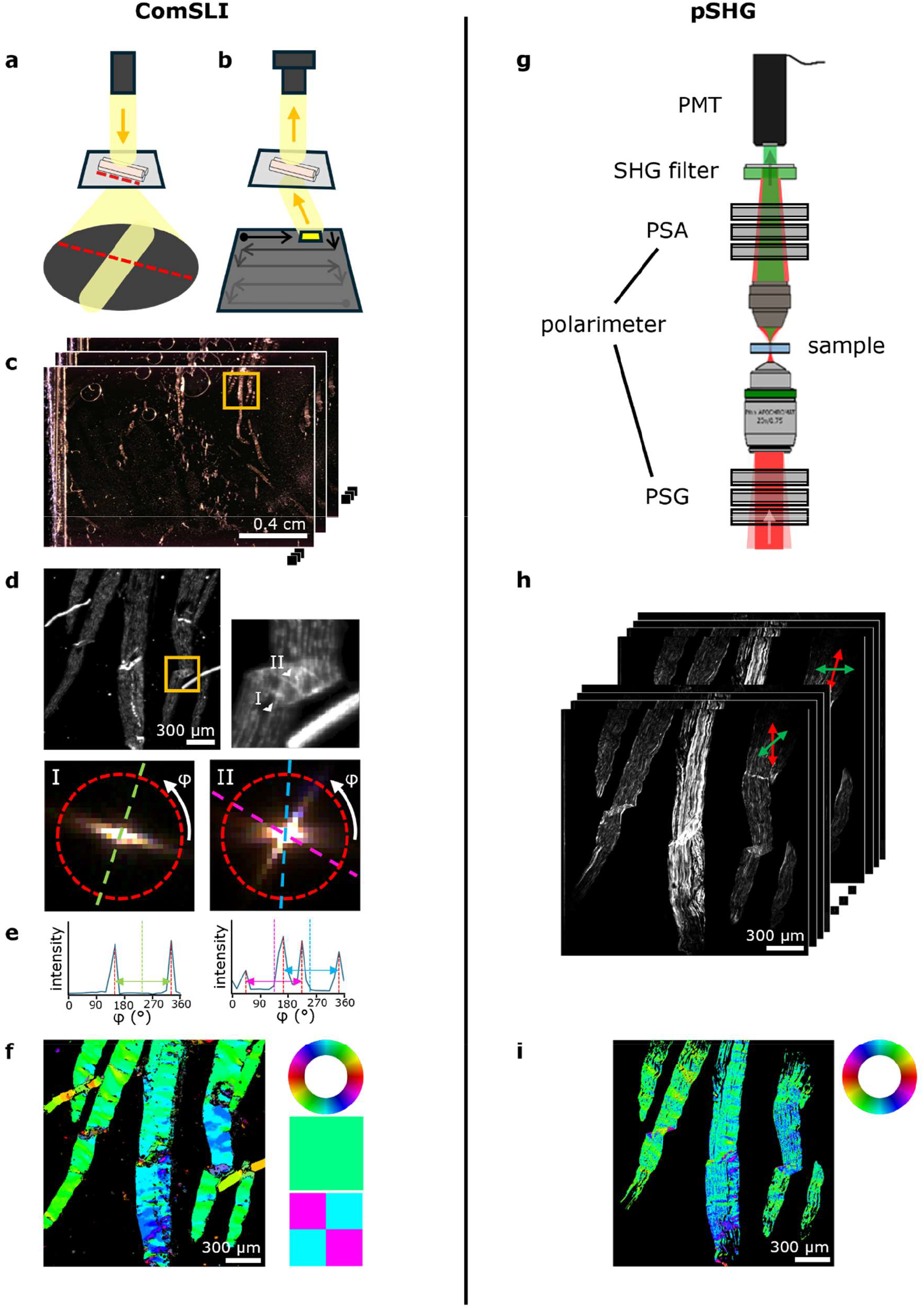
Working principles of ComSLI (left) and pSHG (right). (**a**) Diffraction of light by an in-plane fibre bundle. Scattering occurs mainly in the direction perpendicular to the structure. (**b**) Oblique illumination applied by ComSLI. LEDs are lit sequentially in x-and y-direction as illustrated by the arrows. (**c**) Image series resulting from measurements taken with different LEDs (2.87 μm pixel size, 3.91-4.38 μm lateral resolution). (**d**) Average scattering intensity image of the region indicated by the orange rectangle in (c), showing tendon sections containing both unidirectional fibres and crossing fibres in regions with folded tendon (zoom-in on the top right). Bottom left: scattering pattern for a single pixel containing unidirectional fibres; a single bright line is visible, indicating the orientation of fibres running perpendicular to it (green dashed line). Bottom right: scattering pattern for a single pixel containing overlaid crossing fibres; two bright lines are visible, indicating the orientations of two crossing fibre bundles (blue and pink dashed lines). The locations of the two pixels are indicated by the arrowheads (I) and (II) in the top right image. (**e**) Corresponding line profiles after measuring the intensity along a circle in 15° steps, see red dashed circles in (d). Plotting the intensity against the azimuth angle reveals angles of high scattering intensity (red vertical dashed lines). Taking the value in the centre of the peak pair indicates the fibre orientation (left: green vertical dashed line; right: pink and blue vertical dashed lines). One peak pair indicates the presence of one fibre bundle (left), two peak pairs indicate two crossing fibre bundles (right). (**f**) Fibre orientation map in which the in-plane fibre orientation is indicated by colour, see colour wheel at the top right. Unidirectional fibre orientations are assigned one colour (middle right), crossing fibre orientations are assigned a multi-coloured pixel (bottom right). (**g**) Part of the pSHG setup: the polarisation state generator (PSG) determines the incoming polarisation states of the fundamental beam (red) and the polarisation state analyser (PSA) defines the polarisation of the outgoing SHG beam (green), which is detected by a photomultiplier tube (PMT). (**h**) Image stack obtained from measurements performed with different polarisations on the same tendon sections shown in (d). Here, only linear polarisations are shown (red arrows for incoming and green arrows for outgoing linear polarisations). (**i**) Fibre orientation map showing the in-plane fibre orientations that were retrieved with DSP SHG microscopy (2.37 μm pixel size, 4.6 μm lateral resolution).

For reconstructing the collagen network with pSHG, a series of measurements is being performed with polarised light from a femtosecond pulsed laser source. During the measurement, the incoming polarisation state is being varied with a polarisation state generator (PSG) as shown in Fig. 1g. At each incoming polarisation, the outgoing intensity is being measured for various polarisation states defined by the polarisation state analyser (PSA). This results in a series of intensity images (see Fig. 1h). A combination of linear and circular polarisations is used for double Stokes polarimetry (DSP) to calculate the unidirectional fibre orientation and effective orientation of crossing fibres, while the polarisation-in, polarisation-out (PIPO) method uses only linear polarisation states to determine a single fibre orientation and individual orientations of crossing fibres (see Methods for further details).

Although based on different principles, both ComSLI and pSHG enable determination of collagen fibre orientation. Figures 2a-e show the comparison for the unstained rat tail tendon sample (region marked in Fig. 1c). The results for two other regions within the same sample are shown in Supplementary Fig. 1. Similarity between in-plane fibre orientations determined with both techniques can be appreciated by comparing the FOMs (Fig. 2c).

**Fig. 2.**
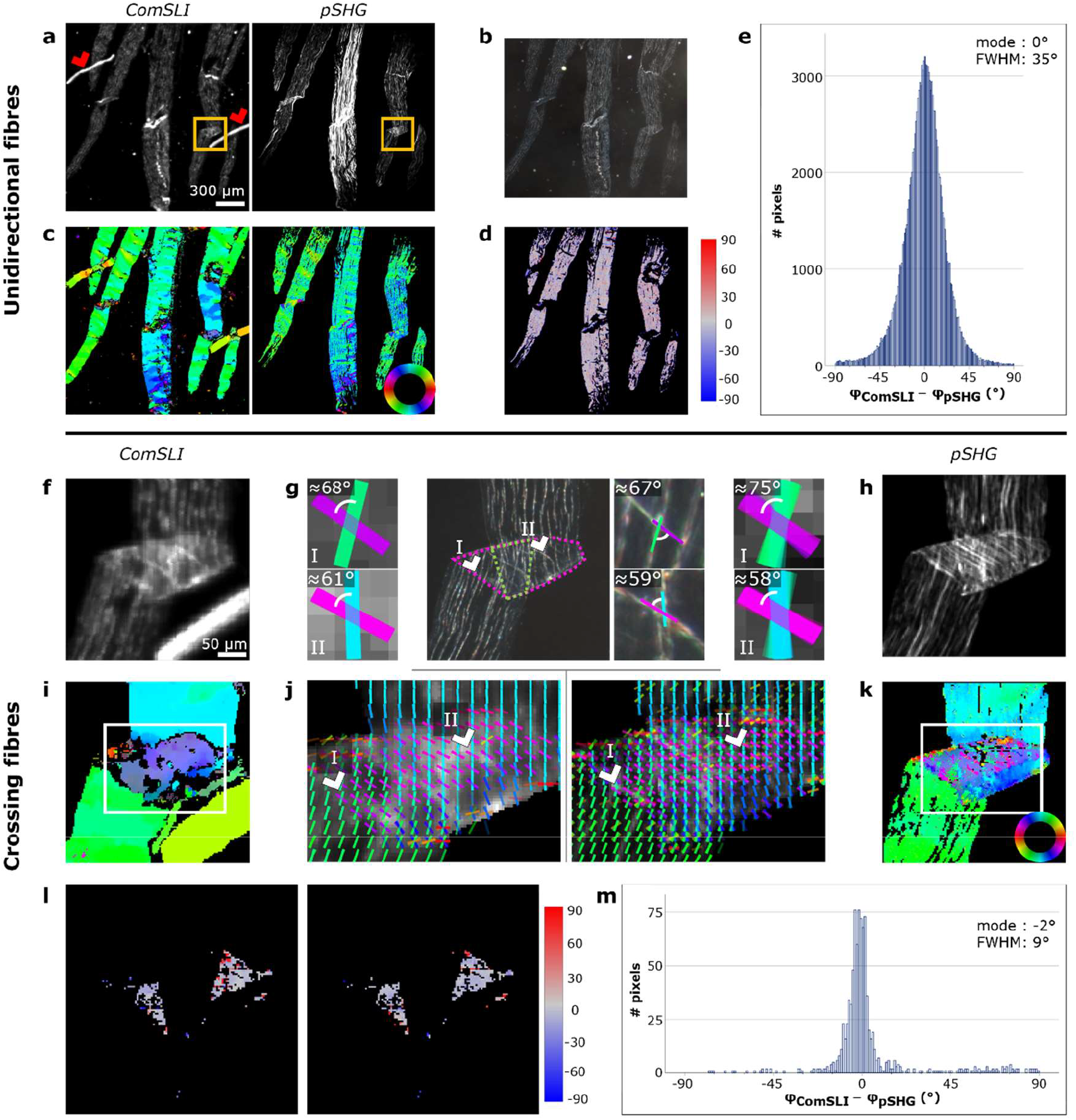
Comparison between ComSLI and pSHG on the in-plane fibre orientations obtained from unidirectional and crossing fibres in unstained rat tail tendon sections. (**a**) Left: Average scattering intensity map from ComSLI (region marked in Fig. 1c). Dust particles are indicated by red arrowheads. Right: Corresponding intensity map from pSHG. (**b**) Corresponding dark-field image. (**c**) FOMs of ComSLI (left) and pSHG (right) obtained with the DSP method; in-plane fibre orientations are indicated by different colours, see colour wheel. (**d**) Difference between unidirectional fibre orientations obtained from ComSLI and pSHG-DSP (in degrees). (**e**) Corresponding histogram. (**f**) Average scattering intensity map from ComSLI for a region with crossing fibres (location indicated by orange rectangles in (a)). (**g**) Middle: Dark-field image. The area containing multiple crossing tendon layers has been indicated by pink dotted lines. Within the region surrounded by green dotted lines, three instead of two crossing fibre layers are present. For two selected fibre crossings, the approximate angle between individual fibre orientations has been indicated in zoom-ins on the right and the individual orientations are colour-coded according to the colour wheel in (k). Left, Right: Distributions of fibre orientations obtained from ComSLI (left) and pSHG-PIPO (right) for regions I (top) and II (bottom) indicated by the arrowheads in (j). Fibre orientations are displayed as coloured bars, and the orientations of 5×5 (ComSLI) and 4×4 (pSHG) pixels are shown on top of each other. (**h**) Intensity map from pSHG. (**i**) Fibre orientation map of ComSLI. (**j**) Distribution fibre orientation maps for the areas indicated in (i) and (k) for ComSLI (left; vectors from 5×5 pixels are shown on top of each other) and pSHG-PIPO (right; vectors from 4×4 pixels are shown on top of each other), respectively. For better visualisation, the signal from the dust particle in the ComSLI measurement has been excluded. (**k**) Fibre orientation map of pSHG-PIPO. (**l**) Difference between fibre orientations obtained from ComSLI and pSHG-PIPO, evaluated only for pixels that yield two different fibre orientations with both techniques and are located within the areas with two crossing fibre layers (as indicated in (g)). Differences are shown separately for one and the other fibre orientation in degrees: the orientation differences were taken between the smallest ComSLI and pSHG values within the defined range (0 - 180°) (left) and between the largest values (right). (**m**) Corresponding histogram including the difference values from both maps in (l).

However, ComSLI assigns orientations to more pixels than pSHG, which is in line with the higher number of pixels showing scattering intensity compared to pSHG intensity (Fig. 2a). This shows the high sensitivity of ComSLI: signal is not exclusively retrieved from collagen, but also from elastin and other fibres, as well as structures like dust particles (indicated by the red arrowheads in Fig. 2a). Examining the differences between the orientations of unidirectional fibres determined with both techniques in a quantitative manner, visualised both in the difference map and in the histogram in Fig. 2d and 2e, respectively, a high correspondence is observed in pixels with a single fibre orientation. Given the non-normal distribution of the data, the mode value and the full-width-at-half-maximum (FWHM) were computed. For this region, the mode was found to be 0° and the FWHM around 35°. The variation in the fibre orientations can be attributed to the different principles employed by both techniques for retrieving the orientation of (collagen) fibres.

Figure 2 panels f-m show the results of analysing a smaller region of the area presented in Fig. 2a- d (see orange rectangles in Fig. 2a) that contains crossing fibres due to folding of the tendon section. The area with fibre crossings is visible in the (average) intensity maps (Fig. 2f, h) and the dark-field image (Fig. 2g), and has been indicated by the pink dotted lines in Fig. 2g (middle). An overall correct distinction between areas with unidirectional fibres and crossing fibres is observed for ComSLI: in the fibre orientation map (Fig. 2i), pixels within the crossing fibres region appear grey-blueish due to the presence of multi-coloured pixels, while pixels outside the crossing fibres region are assigned only one orientation. The distribution of orientations (Fig. 2j, left), shown for the area indicated in Fig. 2i, shows an overall alignment of the orientations with the fibre trajectories expected from the dark-field image. For two locations, this has been highlighted in Fig. 2g where the colour-coded orientations and crossing angles correspond very well with the orientations determined visually from the dark-field image (middle). Within the fold, an area with three instead of two layers of tendon is present (indicated by the green dotted lines in Fig. 2g, middle). ComSLI, however, mostly assigns two orientations to the pixels in this region. The failure to distinguish three fibre orientations can be attributed to the relatively small angle of about 20° between two of the three fibre layers. Depending on the signal-to-noise ratio, the scattering signals of two fibre bundles crossing at an angle smaller than 25° might overlap, yielding only one average fibre orientation. This is also visible in the scattering pattern in Fig. 1d (right), where the horizontal line of high intensity is wider than the other line in the pattern due to the presence of two fibres running in a similar direction.

The orientations of crossing fibres calculated with pSHG-PIPO are similar to the ones obtained with ComSLI (Fig. 2g, j right, k). In regions with two crossing tendon layers, the pSHG-PIPO fibre orientations correspond very well to the visually determined orientations of the dark-field image (Fig. 2g right). In the region with three tendon layers, pSHG-PIPO determines only two orientations as the method for retrieving crossing fibre orientations has been developed for two crossing fibre directions (see Methods). For this reason, only the regions with two tendon layers were included in the quantitative analysis (Fig. 2l, m). Results of the comparison are similar to the ones obtained with unidirectional fibres using the pSHG-DSP method. The lower FWHM value for crossing fibres can be attributed to different pSHG methods. DSP was used to determine the orientation of unidirectional fibres and the effective orientation of crossing fibres. The individual orientations of crossing fibres were determined with the PIPO method. PIPO involves fitting the polarimetric data, while DSP provides direct calculation of the effective orientation from the measured intensity values and, therefore, DSP is more susceptible to noise.

### The influence of tissue staining on orientation retrieval

In order to investigate the influence of the sample preparation process on the determination of collagen orientations, measurements were performed on two differently stained samples. In Fig. 3, the results of the comparison are shown for an eosin-stained sample and an H&E-stained sample. Other regions of interest have been analysed for both samples, the results of which can be found in Supplementary Figs. 2 and 3. Despite the presence of tissue staining, both techniques acquire a decent signal, as can be seen in the (average) intensity maps (Fig. 3a, f). The orientations that are determined from the measured intensities are overall similar for ComSLI and pSHG-DSP, both for the eosinand the H&E-stained sample (Fig. 3c, h). An exception to this can be observed in the regions with crossing fibre bundles (Fig. 3a, b, f, g), where pSHG calculated with DSP retrieves the effective orientation of crossing fibres, while ComSLI shows two orientations per pixel. For discrete orientations of crossing fibres, the pSHG-PIPO method can be applied, similar as for the unstained sample (Fig. 2). We focus further on the orientations of unidirectional fibres determined with the pSHG-DSP method. Quantitatively, differences in the orientation of unidirectional fibres are distributed similarly for both stains, with the same mode value of 1° and FWHMs of about 25° and 23° for eosin and H&E, respectively (Fig. 3e, j). Considering all regions analysed (Fig. 3e, j and Supplementary Figs. 2 and 3), an average mode of 1.7° and FWHM of about 27° were found for eosin and an average mode of 0.3° and FWHM of about 23° were found for H&E. Compared to these parameters determined for the unstained sample (average mode = -0.7°, FWHM ≈ 33°; Fig. 2e, Supplementary Fig. 1e, j), similar mode values and even more narrow distributions were found for the stained samples.

**Fig. 3.**
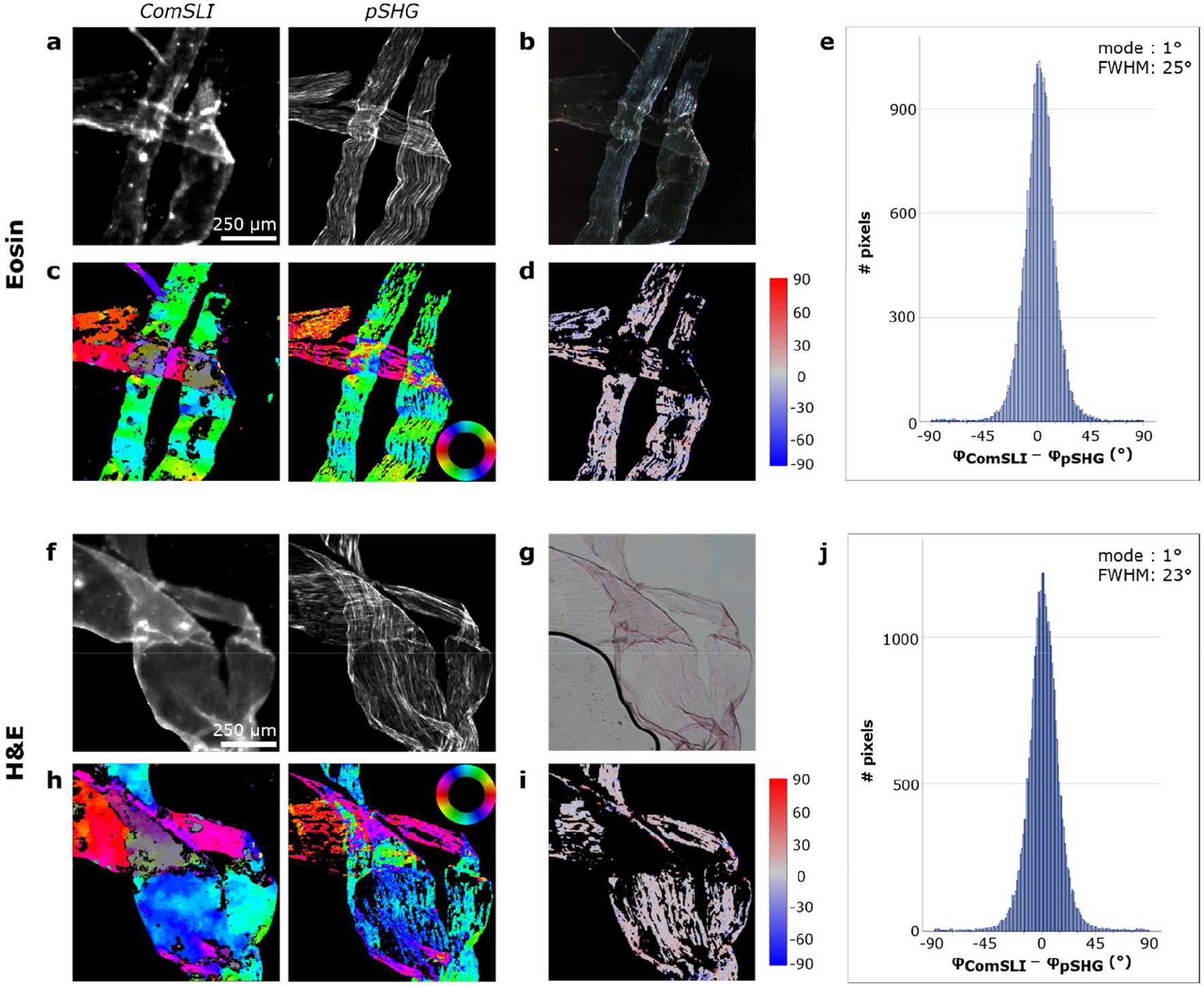
Comparison between ComSLI and pSHG-DSP on the in-plane fibre orientations obtained from unidirectional fibres in differently stained rat tail tendon sections. **(a)** Left: Average scattering intensity map from ComSLI for an eosin-stained sample. Right: Corresponding intensity map from pSHG. (b) Corresponding dark-field image. **(c)** FOMs of ComSLI (left) and pSHG-DSP (right); in-plane fibre orientations are indicated by different colours, see colour wheel. **(d)** Difference between unidirectional fibre orientations from ComSLI and pSHG-DSP (in degrees). **(e)** Corresponding histogram. **(f)** Left: Average scattering intensity map from ComSLI for an H&E-stained sample. Right: Corresponding intensity map from pSHG. **(g)** Corresponding bright-field image. **(h)** FOMs of ComSLI (left) and pSHG-DSP (right); in-plane fibre orientations are indicated by different colours, see colour wheel. **(i)** Difference between unidirectional fibre orientations from ComSLI and pSHG-DSP (in degrees). **(j)** Corresponding histogram.

### The in-plane orientation of out-of-plane tilted fibres

To study the performance on samples with fibres tilted out-of-plane, measurements were performed on rat tail tendon cut under oblique angles of 60° and 90°. While the previous results showed well-aligned collagen fibre bundles in regions with unidirectional fibres, numerous different orientations are observed in the obliquely cut sections (Fig. 4c, h and Supplementary Figs. 4, 5,6). Despite the in-plane organisation being less structured in these samples, both techniques yield similar in-plane fibre orientations, as can be seen from the qualitative comparison of the corresponding FOMs. The quantitative comparisons shown in Fig. 4d, e, i, and j provide the mode values and FWHMs, which were found to be 1° and 27° for the 60°-cut sample, respectively, and 0° and 23° for the 90°-cut sample, respectively. These values are comparable to those found for the samples with in-plane fibres, however, larger differences in orientation are relatively more common among the obliquely cut samples.

**Fig. 4.**
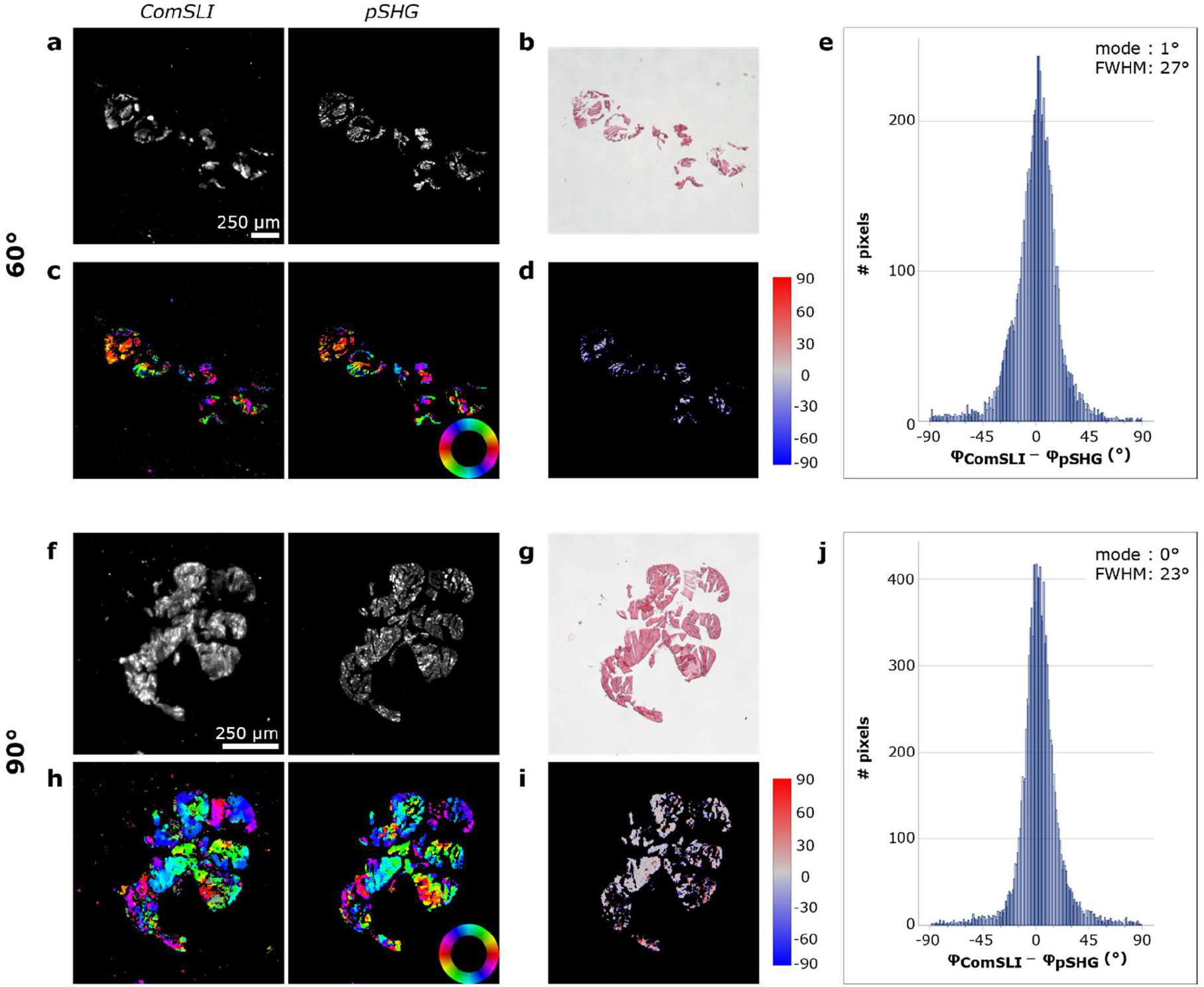
Comparison between ComSLI and pSHG-DSP on the in-plane fibre orientations obtained from obliquely cut rat tail tendon samples (H&E-stained). (**a**) Left: Average scattering intensity map from ComSLI for a 60°-cut tail section. Right: Corresponding intensity map from pSHG. (**b**) Corresponding bright-field image. (**c**) FOMs of ComSLI (left) and pSHG-DSP (right); in-plane fibre orientations are indicated by different colours, see colour wheel. (**d**) Difference between unidirectional fibre orientations from ComSLI and pSHG-DSP (in degrees). (**e**) Corresponding histogram. (**f**) Left: Average scattering intensity map from ComSLI for a 90°-cut tail section. Right: Corresponding intensity map from pSHG. (**g**) Corresponding bright-field image. (**h**) FOMs of ComSLI (left) and pSHG-DSP (right); in-plane fibre orientations are indicated by different colours, see colour wheel. (**i**) Difference between unidirectional fibre orientations from ComSLI and pSHG-DSP (in degrees). (**j**) Corresponding histogram.

### Collagen fibres in bone

For the rat bone section shown in Fig. 5a, two different regions of interest were analysed. The FOMs of those regions again show an overlap in the in-plane fibre orientations determined with both techniques (Fig. 5c). Region I originates from a location containing more longitudinally cut fibres compared to region II, which contains transversely cut fibres. The high amount of small fragments in the pSHG-DSP orientation map of region II resulted in difficulties during the registration of the ComSLI data to the pSHG-DSP data, and the analysis of region II was therefore performed by registering the pSHG-DSP FOM onto the ComSLI FOM (see Methods). The quantitative comparisons of both regions show the same behaviour as the tendon samples (Fig. 5d, e; (I) mode = 2°, FWHM ≈ 22°; (II) mode = 4°, FWHM ≈ 25°). The distribution of the differences is less smooth for region II compared to the distributions obtained for the regions of interest with unidirectional in-plane fibres; similar to the observation on the tendon sections with out-of-plane fibres.

**Fig. 5.**
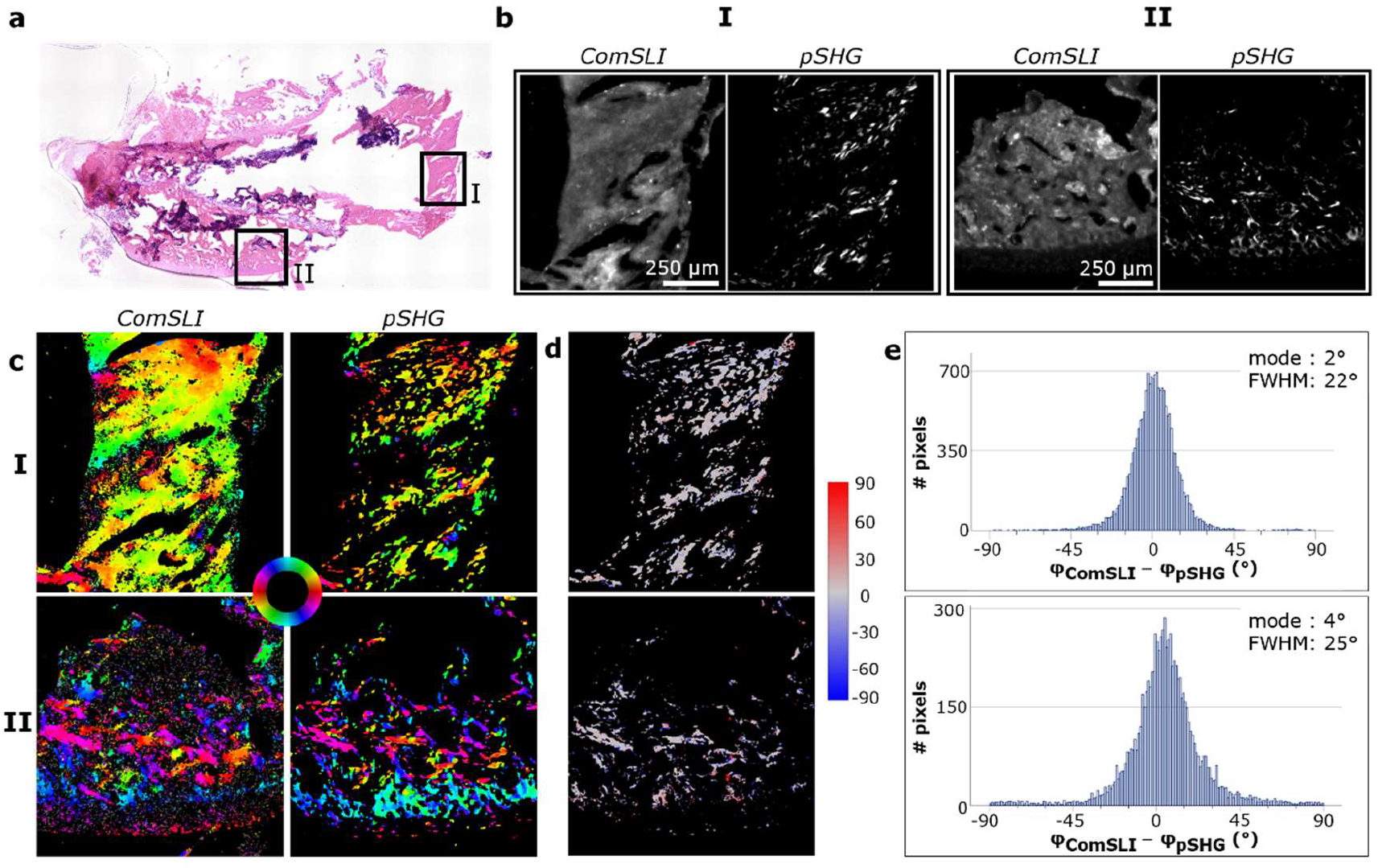
Comparison between ComSLI and pSHG-DSP on the in-plane fibre orientations obtained from unidirectional fibres in a rat bone section (H&E stained). **(a)** Bright-field image; regions measured with both techniques have been indicated with black squares. **(b)** Average scattering intensity map (ComSLI) and intensity map (pSHG) for regions I and II. **(c)** Fibre orientation maps of region I (top) and II (bottom) obtained with ComSLI (left) and pSHG-DSP (right); in-plane fibre orientations are indicated by different colours, see colour wheel. **(d)** Difference between unidirectional fibre orientations from ComSLI and pSHG-DSP (in degrees) for locations I (top) and II (bottom). **(e)** Corresponding histograms for locations I (top) and II (bottom).

## Discussion

The organisation of collagen fibres, structures abundant in numerous types of tissues, largely determines tissue properties. Understanding the relation between organisation and function is therefore important and this requires imaging techniques able to visualise the orientation of fibres in collagen networks. Here, we validate the utility of ComSLI for this purpose by performing comparative measurements with pSHG on collagen-containing rat tissue sections. We started this study by investigating the orientation of collagen fibres in rat tail tendon sections. Within collagen research, rat tail tendon is an often used sample, among others because of its simple structure^7^: collagen fibres are oriented in the same direction as the axis of the tendon, resulting in an organised network with wellknown fibre orientations. In the tendon samples with in-plane fibres, the orientations of unidirectional fibres match the trajectory of the tendon for both ComSLI and pSHG-DSP, therefore suggesting a correct retrieval of the orientations with both techniques. The fact that this holds for both unstained and differently stained samples shows the wide applicability of both techniques. This is advantageous in two ways: first of all, the techniques can be used in a label-free manner. This allows for measuring tissue in its native form, and could therefore speed up the measurement procedure compared to histopathology procedures that include tissue staining. At the same time, measured orientations are not affected by H&E staining, which facilitates the implementation of the techniques into already existing protocols.

The similarity between fibre orientations determined by ComSLI and pSHG-DSP is also observed for obliquely cut tendon sections with fibres tilted out-of-plane. In these sections, fibres are less well aligned, as can be seen in the FOMs, which show the in-plane projection of the fibres (Fig. 4c, h). Since samples were mounted between slides and coverslips, the glass may have pushed the out-of-plane fibres into different in-plane orientations. This would also explain why in-plane fibre orientations were retrieved for the 90°-cut sample with ComSLI; if fibres had been oriented at 90° with respect to the image plane, no in-plane fibre orientation would have been determined. It should be noted here that the samples were cut manually and that a clear measure for determining the inclination angle was absent. It is therefore expected that the real inclination angles in the obliquely cut samples deviate from the 60° and 90° reported in this study.

In principle, both techniques are capable of estimating out-of-plane inclination angles. In pSHG-DSP and PIPO, the achiral and chiral susceptibility ratios are determined, revealing the out-of-plane tilt angle and fibre polarity^36^. In ComSLI, the scattering pattern and the line profile are expected to change in the presence of inclined fibres^30^. For the samples in this study, we observed a bending of the scattering patterns and a decrease in the distance between peaks in the line profiles as expected (Supplementary Fig. 7).

In contrast to tendon, bone has a more complex structure. In addition to collagen fibres, minerals are present to support the stiffness of bone^33,37^. The crystals are mainly directed along the collagen fibre orientation^33^. The complexity of bone tissue seems to have an effect on the determination of the fibre network. For one of the locations studied, oriented structures are only detected in small fragments which hindered the registration of the ComSLI data to the pSHG-DSP data. The other region of interest contains a more aligned fibre network in which the correspondence between both techniques can be better appreciated (Fig. 5c-I).

Looking at the pixel-based differences obtained from the quantitative analyses, orientations obtained with ComSLI and pSHG-DSP are in best agreement for the in-plane fibres in the tendon samples, independent of the stain. In these cases, the histograms display a smooth distribution with low amounts of outliers. A similarly shaped distribution is seen in the bone sample for the region with in-plane fibres. For the other region, containing out-of-plane fibres, the distribution of the differences is less smooth, despite the similar statistical values. This is also observed in the tendon samples with out-of-plane fibres.

While the results show corresponding unidirectional fibre orientations for both techniques, working principles and technical specifications are different. Starting with the technical specifications, pSHG requires more sophisticated optical components and is therefore more difficult to realise and more expensive compared to ComSLI. Requiring only a rotating LED light source or LED display and a camera, ComSLI can be implemented easily and at low costs. With regard to imaging specifications, pSHG is a high-resolution technique^38^, capable of imaging with sub-micron lateral resolution. In this study, a 4x 0.13 numerical aperture (NA) objective was used for pSHG measurements, which gives lower resolution than the maximum possible, to better match the optical lateral resolution of about 4 μm in ComSLI (determined by measuring a USAF resolution target; see Methods). This resolution still allows to distinguish collagen fibres in the tendon sections (10-50 μm)^7^. However, depending on the application, different resolutions may be desired. When a lower resolution is used in pSHG measurements, the microscope’s focal volume can encompass several discrete fibres. The PIPO model of discrete crossing fibres showed consistent results for pSHG with low NA objectives (see Methods). On the other hand, the DSP method provided effective (average) orientations of the crossing fibres applicable for quick assessment of large sample areas. In ComSLI, higher resolutions could be acquired by using an objective with a higher NA, but a trade-off exists between resolution and measurement accuracy. The higher the NA, the larger the acceptance angle, leading to a higher detection rate of light with a non-normal incidence. This affects the signal, and consequently, the determined fibre orientations. Another disadvantage that comes with a higher resolution is a smaller field of view (FoV). While in pSHG the FoV is usually limited to about 1×1 mm^2^, a more than 164 times larger FoV (15.7×10.5 mm^2^) is obtained in ComSLI. In this study, the pSHG FoV was larger due to the lower resolution, but still smaller than the ComSLI FoV (Fig. 1c; the orange rectangle shows the pSHG FoV on top of the ComSLI FoV). Measuring the same area with pSHG requires raster scanning of the sample. With an about three times longer acquisition time, pSHG based on raster scanning is therefore less suited for measuring larger areas and could be replaced with wide-field pSHG microscopy^39^.

In the literature, polarisation-sensitive optical coherence tomography (PS-OCT) is mentioned as a promising technique for visualising collagen networks and overcoming the issue of having a small FoV^10^. The technique is based on the birefringence of collagen, and can be applied *ex-vivo, in-vivo* and in-depth; the last two are currently not possible with ComSLI. However, polarisation-based techniques suffer from a drawback as they struggle to retrieve the individual orientations of crossing fibre bundles^10^. In this study, we performed a comparison between crossing fibre orientations in unstained tendon sections. The retrieval of crossing fibre angles with pSHG requires the acquisition of multiple images with different polarisation states to achieve 3D information that can be compared to the multi-directional results from ComSLI. If only one of the crossing fibres lies within the voxel, this fibre will contribute the most to the measured signal and consequently only one fibre orientation will be determined. In case both fibres lie within the same voxel, the effective (average) fibre orientation is determined with DSP, and discrete orientations of the crossing fibres are determined with the PIPO method. Overall, ComSLI and pSHG-PIPO give similar crossing fibre orientations, which correspond to the actual fibre orientations that are visible in the dark-field image (Fig. 2f-m). In regions with three crossing tendon layers, both techniques fail at resolving all orientations. For pSHG, this can be attributed to the method being developed for distinguishing only two crossing fibre orientations. ComSLI has previously been proven effective in resolving three crossing fibre bundles with crossing angles larger than 25° ^29,40^. Since the section in Fig. 2 contained a 3-layered region with two of the three fibre layers being separated at a smaller angle, only two orientations were resolved.

The comparison shows that ComSLI and pSHG-PIPO give similar orientations when it comes to the retrieval of crossing fibre bundles. In pSHG, optical sectioning provides details on the 3D crossing fibre structures, albeit requiring some input from the user for determining unidirectional and crossing fibre orientations. ComSLI only requires one image processing step: analysing the change in scattering intensity over the different azimuth angles of illumination per pixel. The illumination angles for which the intensity is highest are used for determining the fibre orientations, and these angles depend only on the orientation of the fibres. In this way, fibres crossing at angles larger than 25° can be reliably distinguished from each other. To the best of our knowledge, no other techniques have been demonstrated with which this can be achieved with micrometre resolution and for comparable large fields of view.

The results show differences in pixel density between both techniques. In pSHG-DSP, pixels with unidirectional fibres might be excluded from the analysis due to low signal-to-noise ratio or the polarimetric response not following C6 symmetry. In ComSLI, post-processed signals will be rejected if scattering peaks cannot be matched to a specific fibre orientation due to a noisy signal, however, still more pixels are assigned a value compared to pSHG.

Differences in pixel density can also be attributed to the fact that ComSLI and pSHG rely on different imaging principles, and therefore differ in the origin of the signal that is being measured. ComSLI exploits the anisotropic scattering or diffraction of LED light by directed structures. Influences of the molecular structure underlying different biological tissues have been observed in the scattering patterns, but do not have an effect on the orientation that is determined. All directed structures present within the FoV therefore contribute to the orientations visualised in the FOM, resulting in a high sensitivity, but low specificity in ComSLI. This is different for pSHG which visualises non-centrosymmetric structures such as collagen. Polarimetric SHG therefore has a high specificity for collagen fibres. This additionally explains the pixel density differences observed in our results.

In conclusion, this validation study shows the ability of ComSLI as an imaging technique for retrieving the organisation of collagen fibre networks in biological tissue sections. ComSLI was compared with pSHG, investigating the differences in collagen fibre orientations in rat tendon and bone sections. Overall, there is a clear correspondence in the orientations obtained with both techniques, with the highest degree of similarity found for unidirectional in-plane tendon fibres. The retrieval of orientations is not affected by the presence of H&E stain, and this can therefore simplify the use of the techniques or their implementation in already existing protocols. In addition, both techniques can retrieve the orientations of multi-directional crossing collagen fibres. Differences between ComSLI and pSHG exist, both in the technical specifications and in the principles used for signal retrieval. While raster-scanning pSHG possesses a higher resolution and specificity than ComSLI, the latter has a larger FoV with micrometre resolution and is easier to implement.

## Methods

### Sample preparation

Tendon and bone samples were obtained from 12-week-old albino Wistar rats (State Scientific Research Institute of the Innovative Medical Center, Vilnius, Lithuania). Animals were sacrificed by cervical dislocation. Longitudinally cut rat tail tendon samples (H&E-stained, eosin-stained and unstained), as well as rat trabecular bone samples (H&E-stained) were prepared according to the frozen section method: the samples were placed in a mould with embedding medium and frozen at -40 °C for 35 s. Subsequently, samples were put in the cryotome’s sample holder and sectioned at -24 °C. The tissue slices were then fixed and stained. The obliquely cut (60° and 90°) rat tail tendon samples were prepared by embedding the tissue in paraffin according to a standard protocol, followed by tissue sectioning and staining.

Rat tail tendon samples were sectioned into 5 μm slices, while the bone sample was sectioned into a 10 μm slice (Leica RM2145). H&E-stained and eosin-stained samples were prepared according to a standard H&E staining procedure using Mayer’s hematoxylin and eosin Y. The hematoxylin staining step was omitted for the eosin-stained samples. The animal procedures and experimental protocols were approved by the Lithuanian Animal Care and Use Committee of the State Food and Veterinary Service (Vilnius, Lithuania, approval number G2-156) and were carried out according to the national and European regulations and in compliance with the ARRIVE guidelines.

### Computational scattered light imaging (ComSLI)

Measurements were performed according to the principle shown in Fig. 1. In order to obtain the full scattering patterns shown in this paper (as in Fig. 1d, bottom), measurements were performed with an LED display (Absen LED AW2.8) consisting of 176×176 red, green, and blue (RGB)-LEDs with 2.84 mm pixel pitch. Samples were illuminated using an illumination pattern of 29×29 white LED spots, each consisting of 6×6 adjacent LEDs. For each spot, an image was captured with a CMOS 20 MP RGB camera (BASLER acA5472-17uc USB 3.0), using an exposure time of 10 s, a gain of 2 dB and a gamma of 1. The camera consists of 5472×3648 pixels on a sensor with dimensions of 13.1×8.8 mm^2^. Before reaching the camera, light passed through a lens (Rodenstock Apo-Rodagon-D120 Lens, focal length = 120 mm, focal ratio = 5.6), resulting in a 2.87 μm image pixel size and 15.7×10.5 mm^2^ FoV. The lateral optical resolution was found to be between 3.91 μm and 4.38 μm by measuring a USAF target.

The data for generating the average scattering intensity maps and the FOMs were acquired by performing an angular measurement, i.e., by measuring intensities in a discretised way along a circle of the scattering pattern, therefore reducing measurement time. The bone and tendon samples were illuminated with a torch light (Ledlenser T2 Tactical LED Torch) and with a fibre-coupled LED light source with specifications as described by Georgiadis et al.^32^, respectively. In short, the fibre-coupled LED light source emits white light with a peak wavelength of 443 nm (range 400-750 nm); light is guided through a 2-metre-long multimode fibre, and passes a collimator (FCM1-0.5-CN, Prizmatix) and diffuser (ED1-S20-MD, Thorlabs) for beam shaping before reaching the sample. In all measurements, the sample was illuminated under an approximately 45° angle and the light source was rotated in steps of 15°, resulting in 24 output images per measurement (each of which being the average of 4 images). The exposure time varied between 30 and 100 ms; for each sample individually, a balance between a broad dynamic range and saturation was established. Intensity images were recorded with a camera (BASLER acA5472-17um USB 3.0) with similar specifications as the camera stated above, except of being monochromatic instead of colour-sensitive.

### Polarimetric second harmonic generation (pSHG) microscopy

Polarimetric SHG imaging was performed with a home-built laser scanning microscope. The setup consisted of a 1030 nm femtosecond laser oscillator (FLINT, Light Conversion), a pair of galvanometric mirrors (ScannerMAX, Pangolin Laser Systems), a 4x 0.13 NA excitation objective (Plan Fluor, Nikon), and a 0.4 NA singlet collection lens. A photomultiplier tube (PMT) operating in photon counting mode (H10682-210, Hamamatsu) was used for SHG signal detection. A 750 nm short-pass (Thorlabs) and a 515 nm bandpass (Edmund Optics) filter were placed before the PMT to block the fundamental beam and to collect emitted second harmonic light. To perform polarimetric measurements, a polarisation state generator and analyser (PSG and PSA) were inserted before the excitation objective and before the PMT, respectively (see Fig. 1g). Both the PSG and PSA consisted of a linear polariser, a half-wave plate, and a quarter-wave plate. The PSG components were optimised for 1030 nm, the PSA for 515 nm. To determine the lateral optical resolution, fluorescent spheres were imaged and the FWHM of the smallest spheres observed was determined, resulting in a 4.6 μm lateral optical resolution. The image pixel size was 2.37 μm. FoVs of up to 1.9×1.9 mm^2^ were imaged.

Several methods are available to extract the fibre orientations from pSHG measurements depending on the polarisation states used^35,36,38,41–43^. Here, polarimetric measurements were performed with the DSP method^44^, and crossing fibre areas were analysed with the PIPO method. For DSP SHG microscopy, six PSG states (four linearly and two circularly polarised) were used for excitation. For each PSG state, six PSA states (four linearly and two circularly polarised) were set sequentially to collect polarised SHG signal. For PIPO, eight equally spaced linear incident polarisation states were used. For each incident polarisation state, eight equally spaced outgoing linear polarisation states were used to record SHG intensity images.

### Data processing

#### In-plane fibre orientations

The intensity images acquired with ComSLI were processed with the Scattered Light Imaging ToolboX (SLIX)^45^ to obtain a set of parameter maps. For this study, the average intensity map (showing the average signal intensity across the different angles of illumination) and the FOM (showing the inplane fibre orientations in different colours) were used. Since for each pixel, up to three different fibre orientations can be determined, orientations within one FOM-pixel are represented by 2×2 subpixels. These are single-coloured for one orientation, dual-coloured for two orientations, and consist of three different colours plus a black subpixel for three orientations. In-plane fibre orientations are indicated as azimuth angle and range from 0° to 180°, starting from the horizontal orientation and moving anti-clockwise for larger values. The FOMs were masked with the average intensity map in order to remove background signal with low average scattering signal.

In pSHG-DSP, the in-plane fibre orientations were calculated using the obtained intensity images at various polarization states. Prior to calculations, pixels with an intensity lower than a threshold value (5 counts, compared with the mean of used intensity images) were discarded from further analysis, and intensity images were filtered with a 3×3 median filter.

Using the DSP method, the in-plane average fibre orientation (*δ*) was extracted directly from the polarimetry measurements using equation (1)^44^. In this paper, there is a difference in terminology between ComSLI and pSHG, in order to be consistent with the terminology used in previous papers on both techniques. This means that the in-plane fibre orientation *φ* used in ComSLI, *φ*_ComSLI_, and the comparisons corresponds to the in-plane average fibre orientation in SHG, *δ* = *φ*_pSHG_:

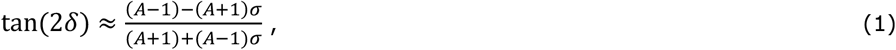

here:

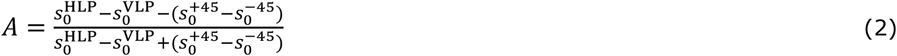

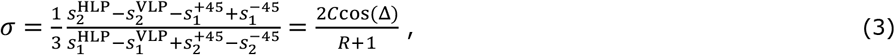

where *C* is the chiral susceptibility tensor element ratio,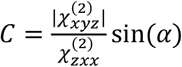, and *α* is the out-of-plane fibre tilt angle. The chiral susceptibility is complex valued, 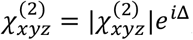, where Δ is the phase difference between the chiral and achiral nonlinear susceptibility tensor components^42^. The *R* is the achiral susceptibility tensor element ratio 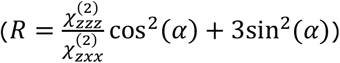. Off resonance conditions are assumed where 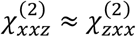 ^42^. The indexes of the tensor elements denote the molecular reference frame, with the *z* axis oriented along the collagen fibre and the *x, y* axes perpendicular to the fibre. The molecular reference frame is related to the laboratory reference frame through matrix rotation.

The Stokes components *s* are calculated from the sample’s response to different linear and circular polarisation states (HLP/VLP: horizontal/vertical linear polarisation, +/-45°: diagonal/anti-diagonal linear polarisation, RCP/LCP: right/left handed circular polarisation). The superscript of *s* denotes the incident polarization prepared by the PSG, and the subscript denotes the outgoing Stokes vector component of the SHG signal determined using the PSA. The *δ* determination range can be expanded to [-π;π] with SHGLD and SHG45 parameters^44^:

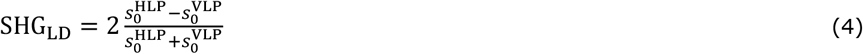

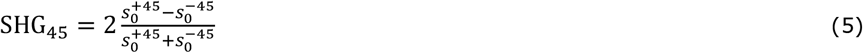

It is not possible to obtain the individual orientations of discrete crossing fibres with DSP, only the effective (average) orientation. Instead, the polarisation-in, polarisation-out (PIPO) pSHG method was used for the area of crossing fibres. In PIPO microscopy, the SHG intensity depends on the orientation of incoming polarisation (*θ*) and outgoing polarisation (*ϕ*) and is proportional to:

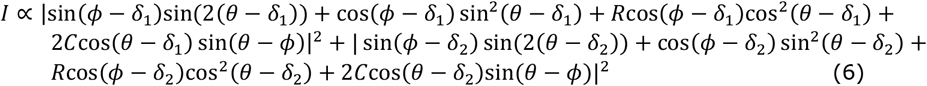

The ultrastructural information of collagen, such as the orientation of two crossing fibres (*δ*_1_ and *δ*_2_) and the susceptibility tensor ratios R and C, can be calculated by fitting the intensity equation (equation (6)) for each pixel of the image set using custom MATLAB software. The R and C ratios were assumed to be the same for both crossing fibres.

#### Image registration

In order to quantitatively compare the results from both techniques, first pre-processing of the inplane orientation maps was performed in Fiji^46^. For this, the ComSLI orientation maps were upscaled to match the pSHG pixel size. Subsequently, the orientation map was registered to the pSHG orientation map by manually applying translation and rotation, after which the orientation map was cropped to match the FoV of pSHG. In addition, the orientation values in the pSHG maps were changed from radians to degrees and adjusted to let the coordinate system match the one used in ComSLI.

One region within the bone sample (region II) consisted of numerous small fragments, preventing us from performing the registration as mentioned above. Instead, the pSHG FOM was registered onto the ComSLI FOM and the applied transformations were used to pre-process the ComSLI orientation maps.

#### Quantitative comparison

Following pre-processing, the set of images was compared in a pixel-wise manner using a self-written Python script. For each pixel, first the number of orientations was determined. In case both techniques did not find an orientation, or they did not find the same amount of orientations, the pixel was excluded from evaluation. In case both techniques retrieved a single fibre orientation, a direct comparison was performed by calculating the difference in degrees between both orientations as follows:

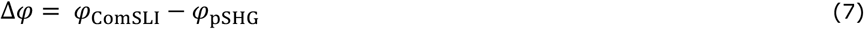

Evaluating pixels with two orientations in both techniques, which was applied only to the unstained in-plane tail section, was done by ordering the orientations from small to large for both techniques and subsequently calculating the difference between the first ComSLI and pSHG orientations and between the second ComSLI and pSHG orientations using the formula above. The differences were visualised in a difference map and histogram, using both Python and Fiji^46^, and IBM SPSS Statistics (28.0.1.0 (142)), respectively.

## Data availability

The datasets generated and analysed during the current study are available from the corresponding authors on reasonable request.

## Acknowledgements

The work was supported by the National Sciences and Engineering Research Council of Canada (NSERC) (RGPIN-2024-06356) and the Lietuvos Mokslo Taryba (LMT) (project no. P-MIP-24-644). L.E. received funding from the Deutsche Forschungsgemeinschaft (DFG) Project no. 498596755. M.A. received funding from the NSERC grant no. QSP-052-1. The authors thank D. Bulotiene, National Cancer Institute, Vilnius, for sample preparations.

## Author contributions

L.E. performed the ComSLI measurements, analysed the ComSLI data, performed the comparison between ComSLI and pSHG, wrote the first version of the manuscript, and prepared the figures. V.M. performed the pSHG measurements, acquired the bright-field and dark-field images, calculated the unidirectional fibre orientations from the pSHG data using DSP, calculated the crossing fibre orientations using PIPO, wrote part of the pSHG methodology and contributed to the editing of the manuscript. M.A. prepared the right side concerning the pSHG methodology of figure 1, wrote part of the pSHG methodology, and contributed to the data interpretation and editing of the manuscript. H.A. contributed to the data interpretation and editing of the manuscript. V.B. contributed to the study design, data interpretation, supervision, and editing of the manuscript. M.M. contributed to the study design, data interpretation, supervision, and editing of the manuscript. All authors reviewed the manuscript.

## Additional Information

**Supplementary Information** accompanies this paper.

## SUPPLEMENTARY INFORMATION

**Supplementary Fig. 1.**
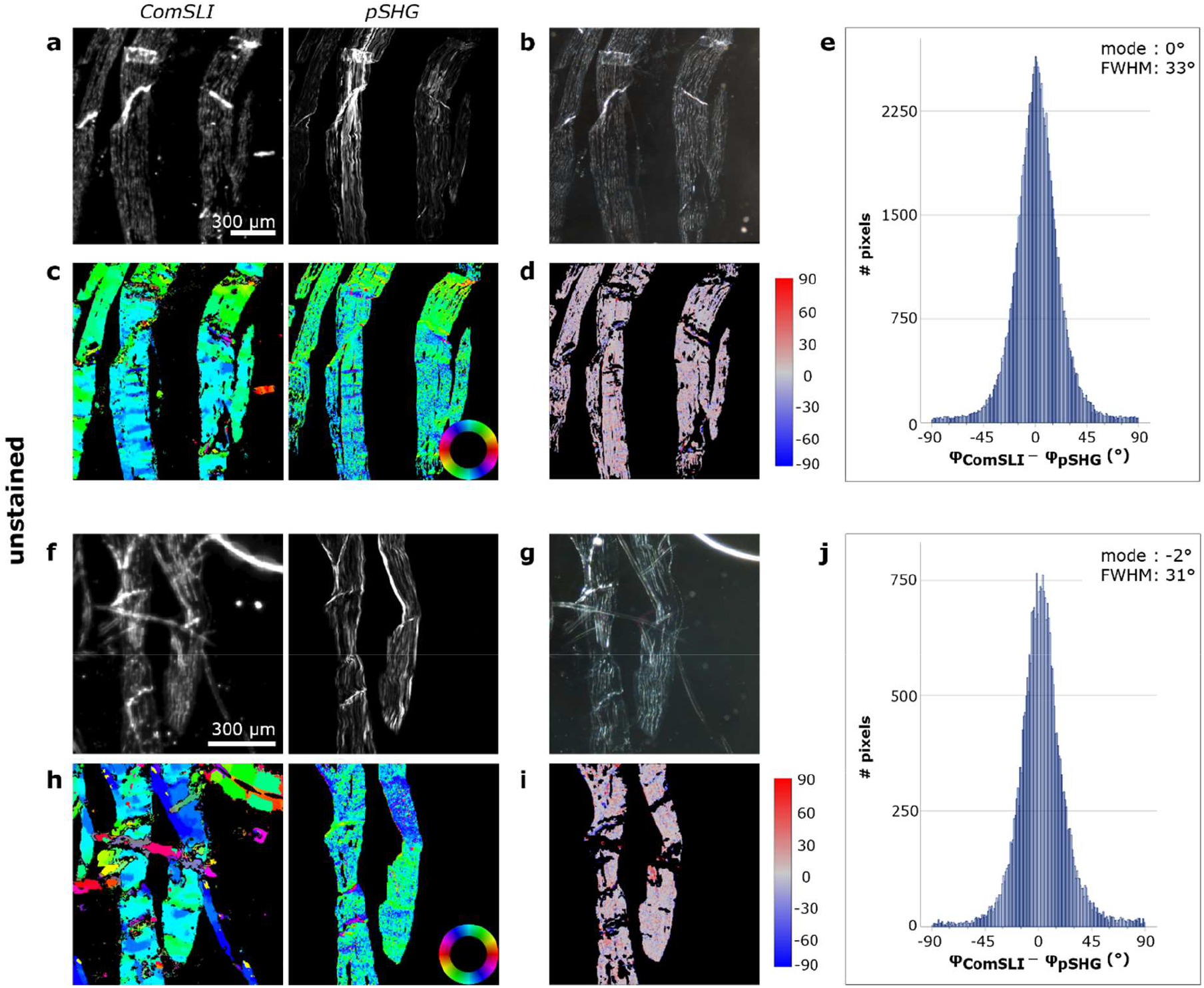
Comparison between ComSLI and pSHG-DSP on the in-plane fibre orientations obtained from unidirectional fibres in unstained rat tail tendon sections. (**a**,**f**) Left: Average scattering intensity maps from ComSLI. Right: Corresponding intensity maps from pSHG. (**b**,**g**) Corresponding dark-field images. (**c**,**h**) In-plane fibre orientation maps of ComSLI (left) and pSHG-DSP (right); orientations are indicated by different colours, see colour wheels. (**d**,**i**) Difference between the unidirectional fibre orientations obtained from ComSLI and pSHG-DSP (in degrees). (**e**,**j**) Corresponding histograms.

**Supplementary Fig. 2.**
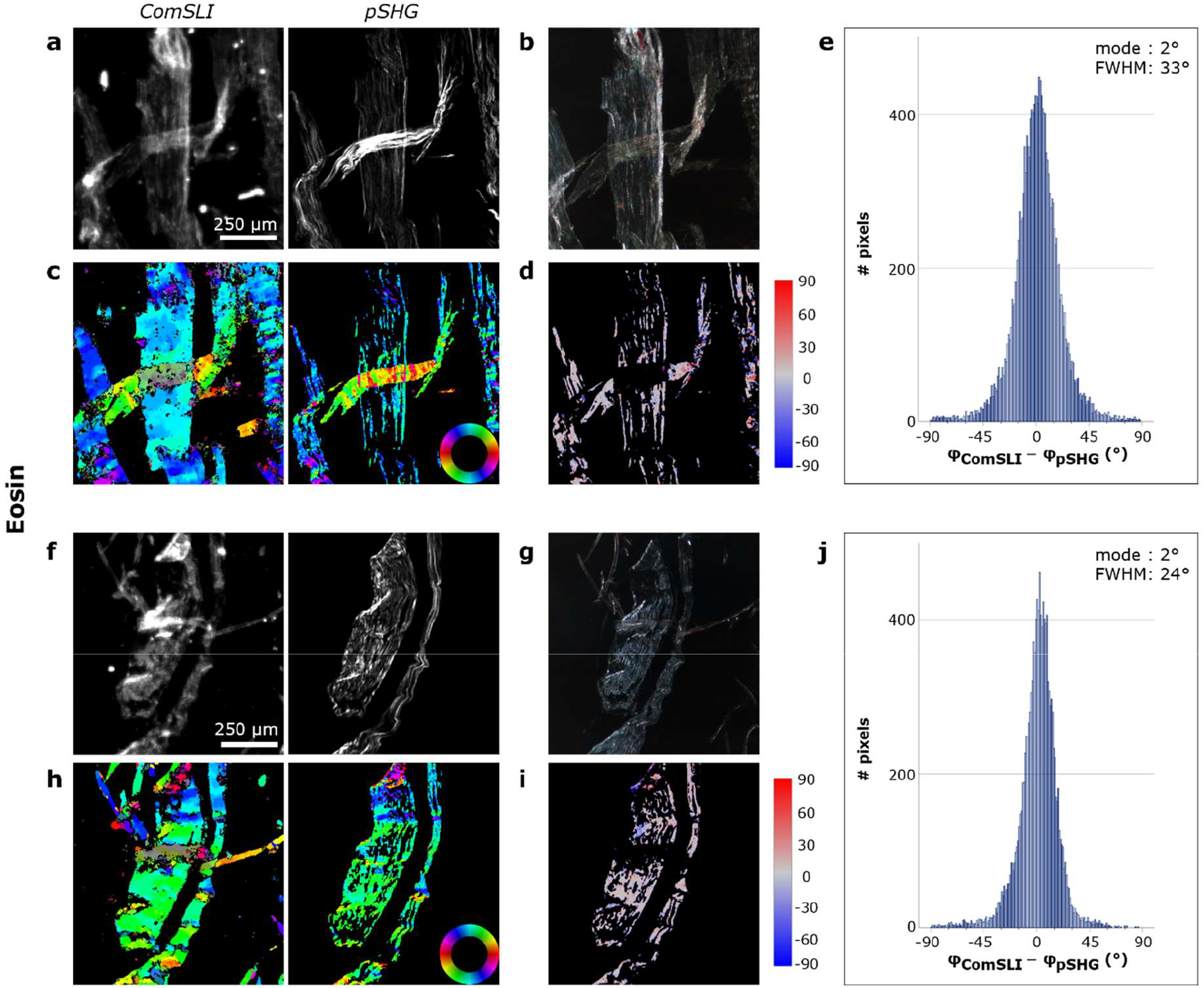
Comparison between ComSLI and pSHG-DSP on the in-plane fibre orientations obtained from unidirectional fibres in eosin-stained rat tail tendon sections. (**a**,**f**) Left: Average scattering intensity maps from ComSLI. Right: Corresponding intensity maps from pSHG. (**b**,**g**) Corresponding dark-field images. (**c**,**h**) In-plane fibre orientation maps of ComSLI (left) and pSHG-DSP (right); orientations are indicated by different colours; see colour wheels. (**d**,**i**) Difference between the unidirectional fibre orientations obtained from ComSLI and pSHG-DSP (in degrees). (**e**,**j**) Corresponding histograms.

**Supplementary Fig. 3.**
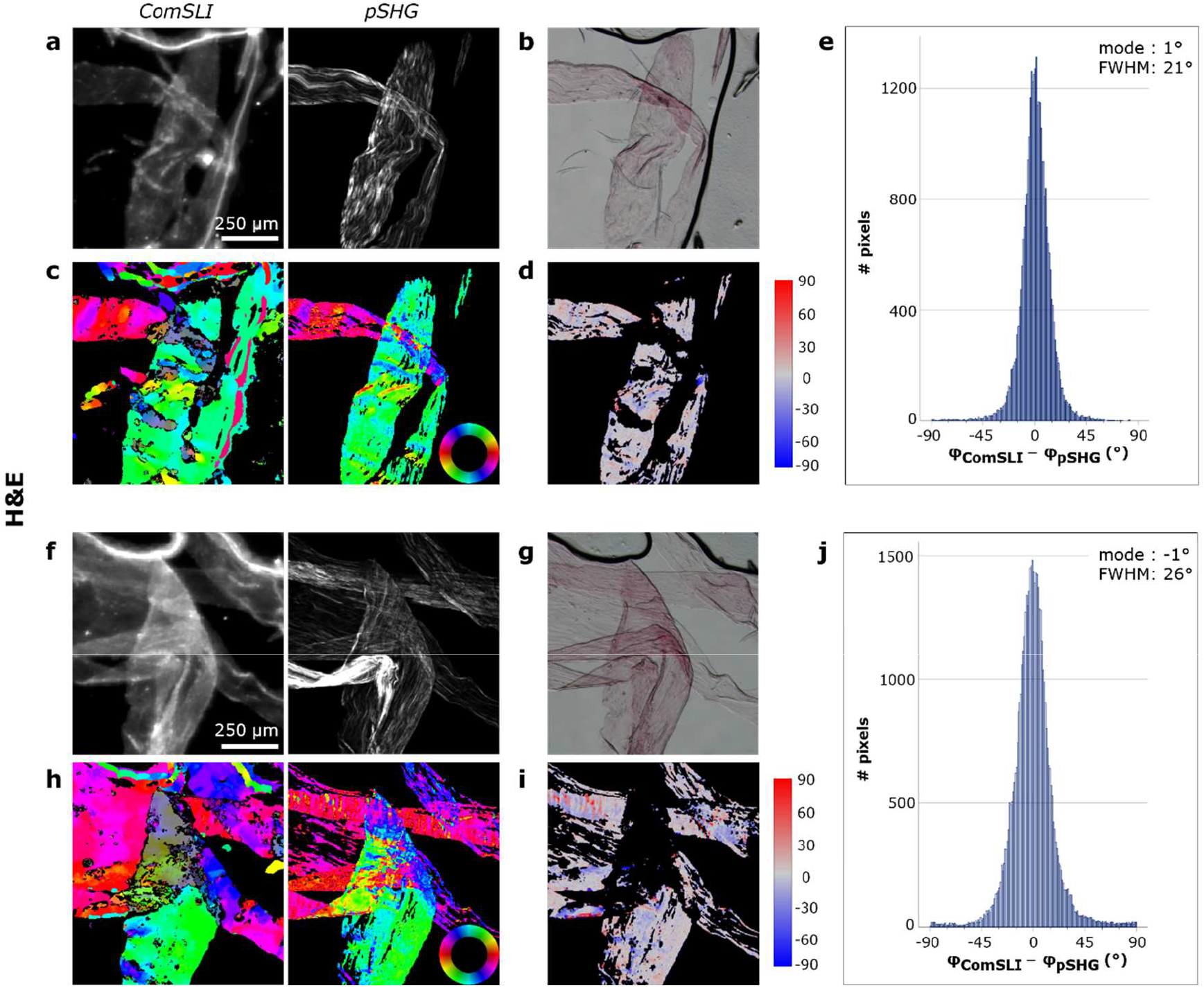
Comparison between ComSLI and pSHG-DSP on the in-plane fibre orientations obtained from unidirectional fibres in in-plane H&E-stained rat tail tendon sections. (**a**,**f**) Left: Average scattering intensity maps from ComSLI. Right: Corresponding intensity maps from pSHG. (**b**,**g**) Corresponding bright-field images. (**c**,**h**) In-plane fibre orientation maps of ComSLI (left) and pSHG-DSP (right); orientations are indicated by different colours; see colour wheels. (**d**,**i**) Difference between the unidirectional fibre orientation obtained from ComSLI and pSHG-DSP (in degrees). (**e**,**j**) Corresponding histograms.

**Supplementary Fig. 4.**
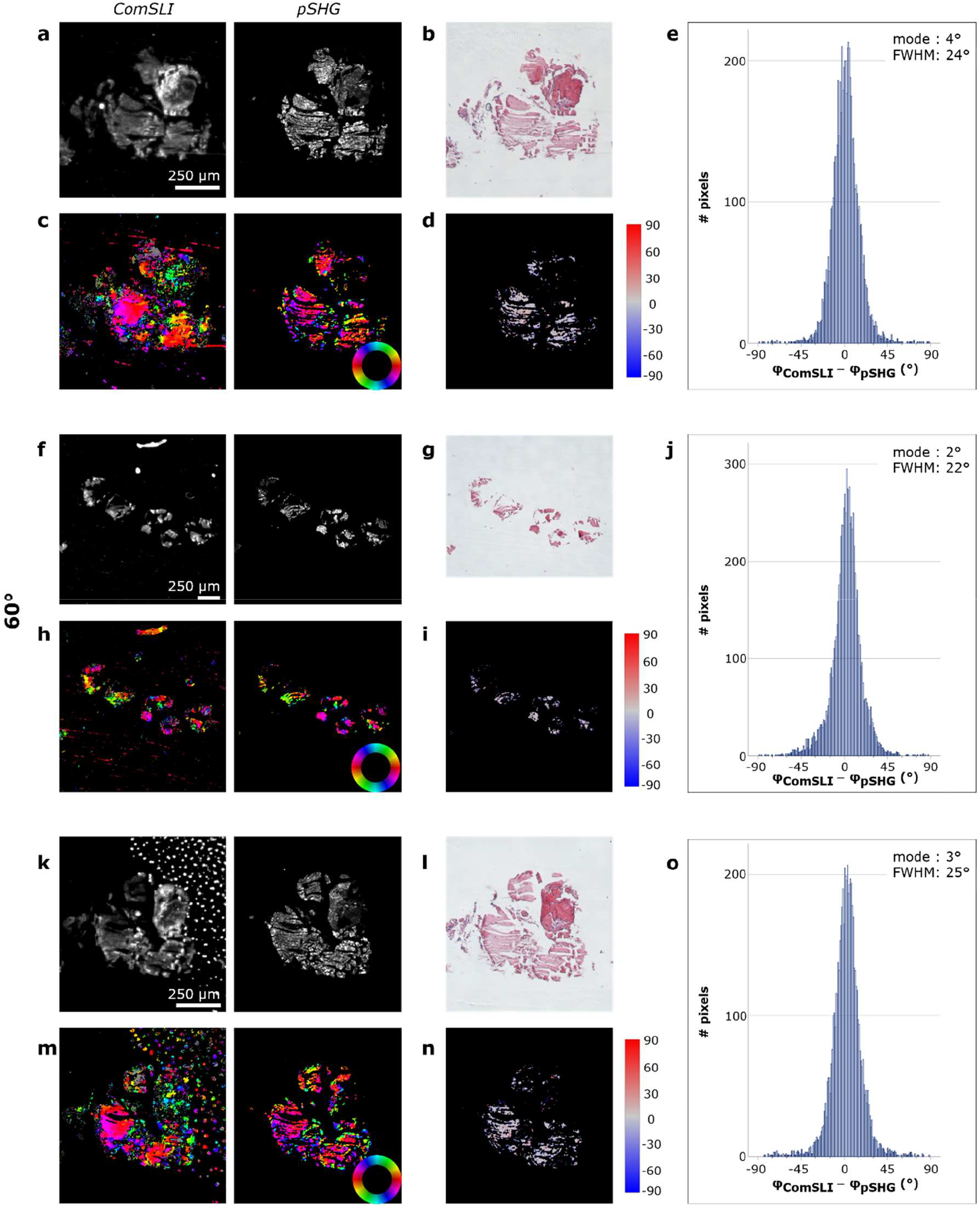
Comparison between ComSLI and pSHG-DSP on the in-plane fibre orientations obtained from 60°-cut rat tail tendon sections (H&E-stained). (**a**,**f**,**k**) Left: Average scattering intensity maps from ComSLI. Right: Corresponding intensity maps from pSHG. (**b**,**g**,**l**) Corresponding bright-field images. (**c**,**h**,**m**) In-plane fibre orientation maps of ComSLI (left) and pSHG-DSP (right); orientations are indicated by different colours; see colour wheels. (**d**,**i**,**n**) Difference between the unidirectional fibre orientations obtained from ComSLI and pSHG-DSP (in degrees). (**e**,**j**,**o**) Corresponding histograms.

**Supplementary Fig. 5.**
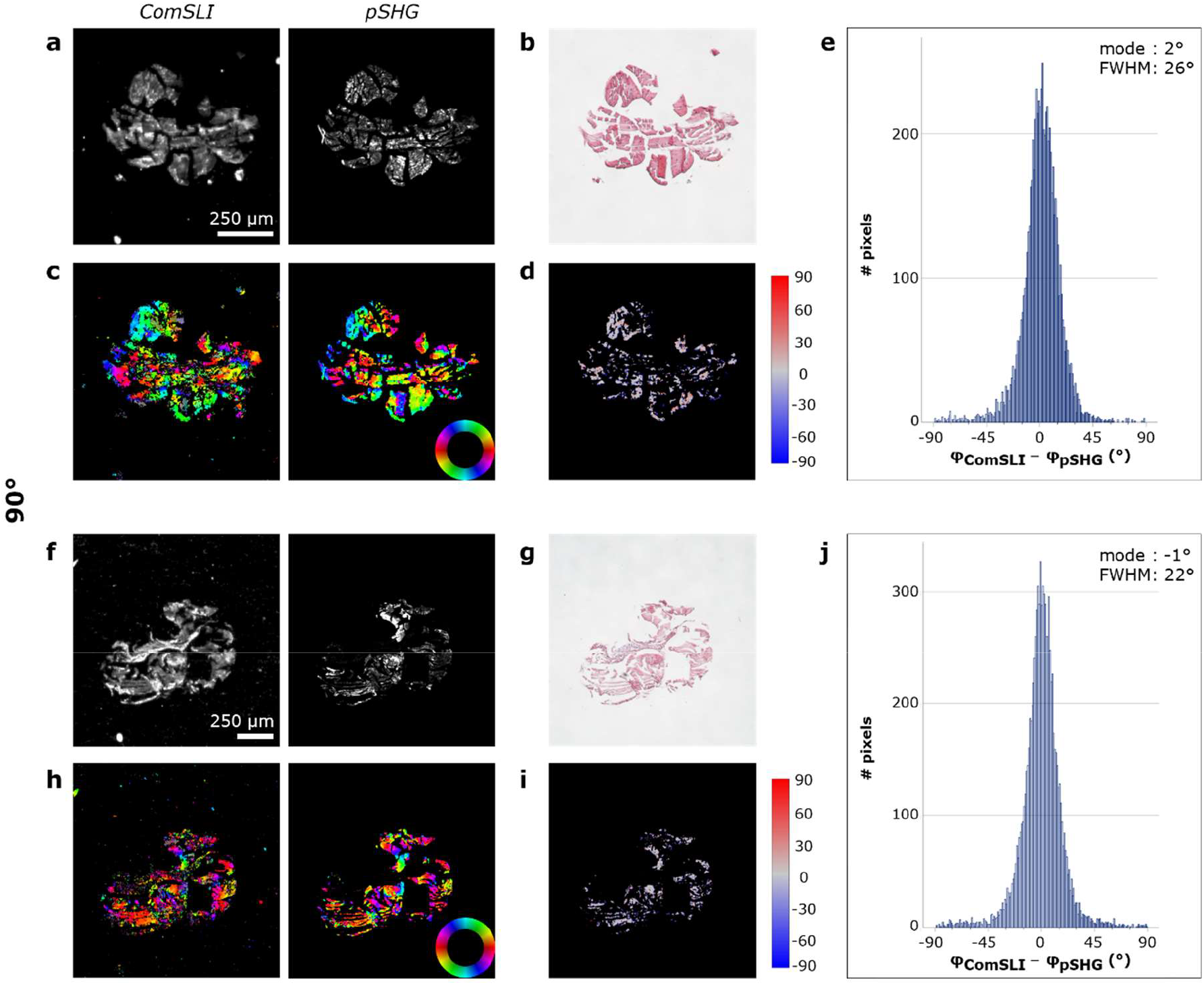
Comparison between ComSLI and pSHG-DSP on the in-plane fibre orientations obtained from 90°-cut rat tail tendon sections (H&E-stained). (**a**,**f**) Left: Average scattering intensity maps from ComSLI. Right: Corresponding intensity maps from pSHG. (**b**,**g**) Corresponding bright-field images. (**c**,**h**) In-plane fibre orientation maps of ComSLI (left) and pSHG-DSP (right); orientations are indicated by different colours; see colour wheels. (**d**,**i**) Difference between the unidirectional fibre orientations obtained from ComSLI and pSHG-DSP (in degrees). (**e**,**j**) Corresponding histograms.

**Supplementary Fig. 6.**
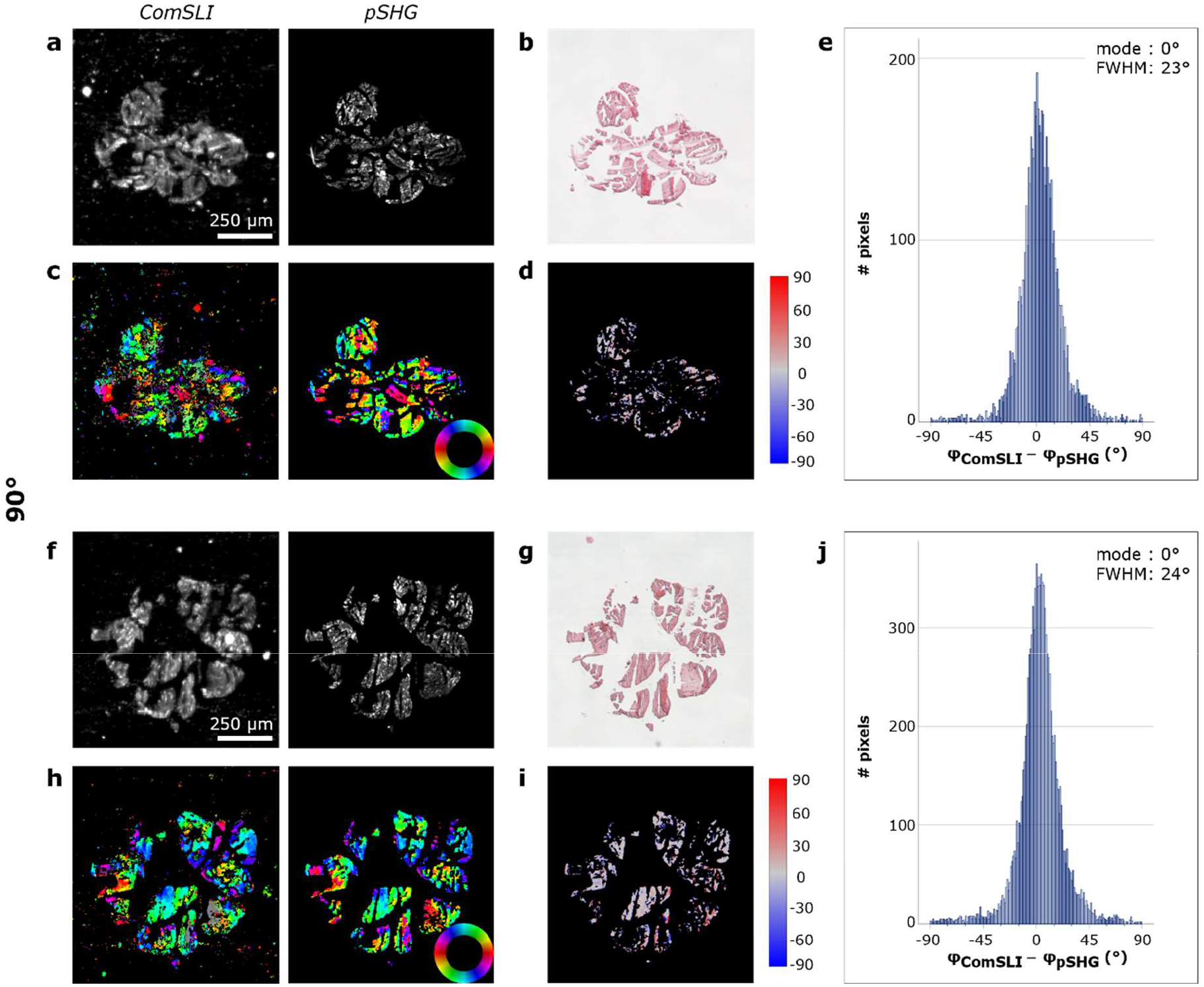
Comparison between ComSLI and pSHG-DSP on the in-plane fibre orientations obtained from 90°-cut rat tail tendon sections (H&E-stained). (**a**,**f**) Left: Average scattering intensity maps from ComSLI. Right: Corresponding intensity maps from pSHG. (**b**,**g**) Corresponding bright-field images. (**c**,**h**) In-plane fibre orientation maps of ComSLI (left) and pSHG-DSP (right); orientations are indicated by different colours; see colour wheels. (**d**,**i**) Difference between the unidirectional fibre orientations obtained from ComSLI and pSHG-DSP (in degrees). (**e**,**j**) Corresponding histograms.

**Supplementary Fig. 7.**
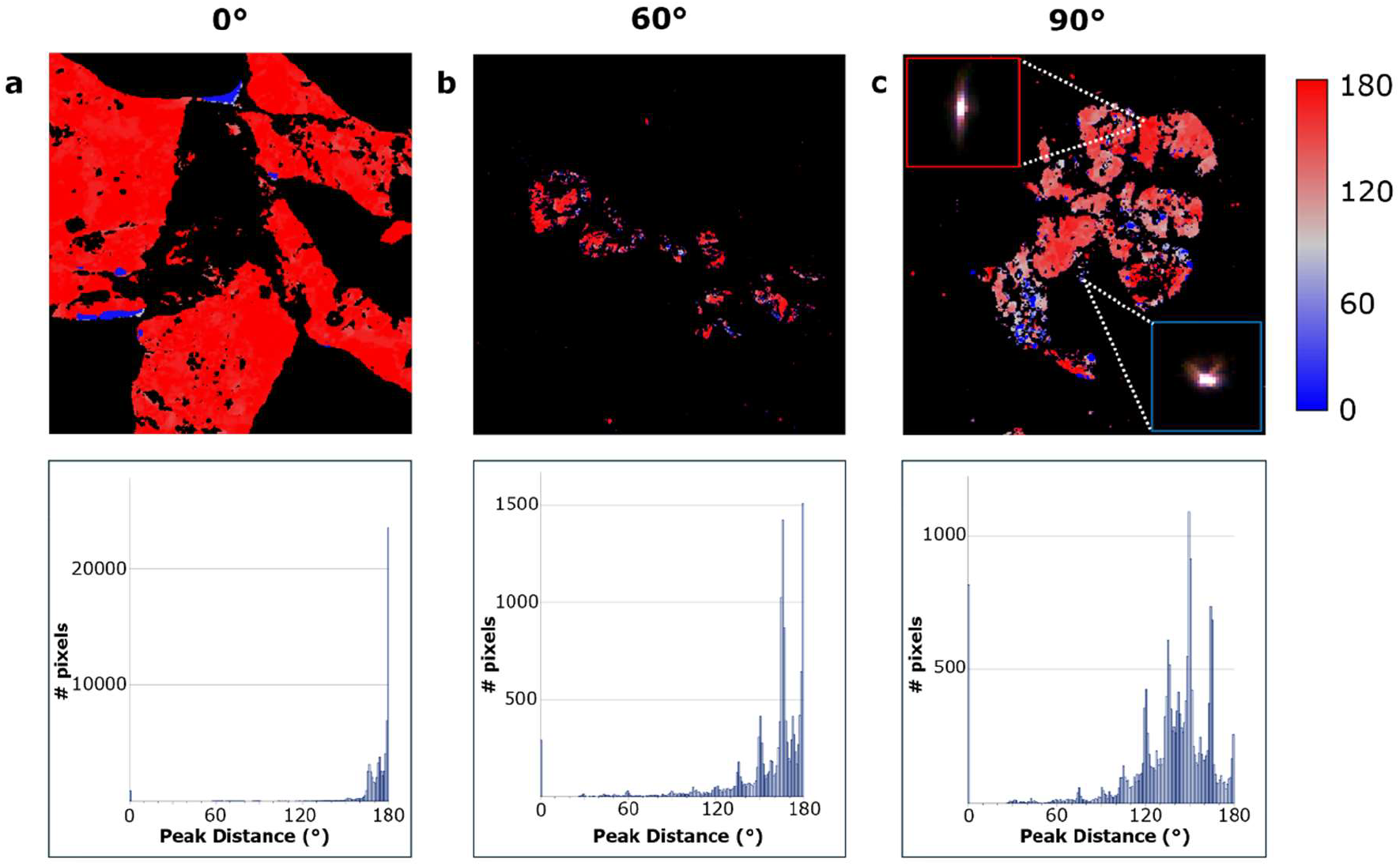
Peak distance maps and scattering patterns for rat tail tendon samples with different fibre inclination angles measured with ComSLI (H&E-stained). (**a**) Peak distance (°) map (top) for H&E-stained tail sections containing mostly in-plane fibre orientations and corresponding histogram (bottom) (mean = 171°); the peak distance map has been masked with the average scattering intensity map and the orientation map in order to get peak distances only for unidirectional collagen fibres. (**b**) Peak distance (°) map (top) for an H&E-stained tail section obliquely cut under 60° and corresponding histogram (bottom) (mean = 155°). (**c**) Peak distance (°) map (top) for an H&E-stained tail section obliquely cut under 90° and corresponding histogram (bottom) (mean = 134°); the inserts show the scattering patterns of two image pixels: one corresponding to a pixel with a larger peak distance (red rectangle) and one corresponding to a pixel with smaller peak distance (blue rectangle), indicating more inclined fibres.

